# Chemosensory ERP Suggest Peripherally Driven Olfactory-Trigeminal Interactions in Healthy Older Adults

**DOI:** 10.64898/2026.08.25.746692

**Authors:** Sarah Brosse, Olivier Fortier-Lebel, Émilie Hudon, Keven Lapointe, Johannes Frasnelli

## Abstract

The olfactory and intranasal trigeminal systems interact closely, influencing chemosensory perception, yet the mechanisms underlying their interaction remain poorly understood and have been studied mainly in young adults. We aimed to characterize olfactory-trigeminal interactions in aging by comparing electrophysiological and behavioral responses under ipsilateral and contralateral olfactory-trigeminal co-stimulation, to determine the relative contributions of peripheral and central mechanisms. Using chemosensory event-related potentials and a localization task, 44 healthy older adults (66.3 ± 4.6 years; 29 women) were tested under four conditions: pure trigeminal (carbon dioxide; CO₂), pure olfactory (2-phenylethanol; PEA), ipsilateral co-stimulation (PEA+CO₂ in the same nostril), and contralateral co-stimulation (PEA+CO₂ in opposite nostrils). Ipsilateral, but not contralateral olfactory-trigeminal co-stimulation, improved trigeminal localization performance and induced larger late positive component amplitudes. Together, these findings suggest that olfactory-trigeminal interactions are driven primarily by peripheral rather than central mechanisms. This study also provides normative CSERP data for healthy older adults.

## Introduction

The olfactory and intranasal trigeminal systems jointly contribute to chemosensory perception (Hummel & Frasnelli, 2019). Whereas the olfactory system detects odorants via stimulation of the olfactory nerve (Cranial Nerve I), the trigeminal system conveys somatosensory information evoked by chemical substances – such as irritation, cooling, warming, and tingling – through activation of the trigeminal nerve (Cranial Nerve V) (Terrier et al., 2022).

Most odorants stimulate both the olfactory and trigeminal systems and are therefore considered mixed olfactory-trigeminal stimuli (Doty et al., 1978). This makes it challenging to isolate the two systems. Nevertheless, certain compounds selectively target one system. For example, vanillin (vanilla odor) and, in low concentrations 2-phenylethanol (PEA; rose odor) are considered pure olfactory stimuli (Doty et al., 1978; Frasnelli et al., 2011). Conversely, carbon dioxide (CO2), which is odorless, is classified as a pure trigeminal stimulus (Albrecht et al., 2010).

Although the olfactory and trigeminal systems are anatomically independent and convey sensations via separate cranial nerves, they are functionally interconnected. In fact, individuals who lose their sense of smell, typically also exhibit reduced trigeminal sensitivity (Hummel et al., 2003) and vice-versa (Husner et al., 2006). In addition, the two systems can mutually suppress and/or enhance each other (Brand, 2006; Cain & Murphy, 1980; Kobal & Hummel, 1988; Livermore et al., 1992; Migneault-Bouchard et al., 2025), indicating that olfactory-trigeminal co-stimulation does not simply reflect the sum of unimodal responses. In line with this, such co-stimulation activates a broader network of brain regions (Boyle, Frasnelli, et al., 2007; Boyle, Heinke, et al., 2007) and yields chemosensory event-related potentials (CSERP) with larger amplitudes and shorter latencies compared to either stimulus alone (Kobal & Hummel, 1988; Livermore et al., 1992).

Understanding this interaction is crucial, as neurodegenerative diseases such as Parkinson’s disease exhibit a disease-specific pattern of chemosensory alteration (Tremblay & Frasnelli, 2021). Interestingly, the intimate connection between olfactory and trigeminal systems is affected in a disease-specific way in Parkinson’s disease. While they typically present with a very early pronounced olfactory decline (Marin et al., 2018), Parkinson’s patients do not exhibit the loss of trigeminal sensitivity usually linked to olfactory loss (Tremblay & Frasnelli, 2021). To comprehend the underlying pathophysiological mechanisms, one must understand the olfactory-trigeminal interaction in health first.

In fact, the neural mechanisms underlying the olfactory-trigeminal interaction remain poorly understood, partly because the two systems can interact both peripherally, i.e., at the level of the mucosa, and centrally, i.e., in the olfactory bulb and on cortical levels (for a review, see Migneault-Bouchard et al., 2025). Most evidence on olfactory-trigeminal interactions comes from studies in which both olfactory and trigeminal stimuli were presented to the same nostril (i.e., ipsilateral co-stimulation), where interactions could occur at either the peripheral and/or central level. To disentangle central from peripheral interactions, one can compare ipsilateral co-stimulation from contralateral co-stimulation where the olfactory stimulus is delivered to one nostril while the trigeminal stimulus is delivered to the other nostril (Cain & Murphy, 1980). If an effect is observed only in the contralateral condition, but not in the ipsilateral, it must be the result of a central interaction, as the stimuli are physically separated on the level of the mucosa.

Importantly, most studies investigating these interactions were carried out in young adults, even though age is one of the factors that influences perception in both sensory systems. While physiological aging is associated with gradual decline in olfactory function (Doty et al., 1984; Doty & Kamath, 2014; Hummel et al., 2007; Oleszkiewicz et al., 2019), intranasal trigeminal function appears to be relatively preserved, showing only a modest age-related reduction (Frasnelli & Hummel, 2003; Hummel et al., 2003; Laska, 2001; Mai, Hernandez, et al., 2025). Together, these findings suggest that aging does not uniformly affect both systems.

To address these gaps, we investigated olfactory-trigeminal interactions in healthy older adults using CSERP. This non-invasive electrophysiological technique provides objective measures of olfactory and trigeminal cortical processing with millisecond precision, enabling the temporal characterization of sensory and cognitive stages that cannot be captured by other neuroimaging methods (Gudziol & Guntinas-Lichius, 2019; Kobal & Hummel, 1988; Osman & Silas, 2015). CSERP are largely independent of response bias, making them a valuable complement to psychophysical measures. They can serve as reliable biomarkers for the assessment of olfactory dysfunction (Baranwal et al., 2024; Kobal & Hummel, 1998; Rombaux et al., 2006). The CSERP waveform typically exhibits a negative deflection around 350-550 ms after stimulus presentation, referred to as the N1, and a positive deflection between approximately 400 and 800 ms, termed the Late Positive Component (LPC) (Pause et al., 1996; Thesen & Murphy, 2002). N1 amplitude and latency are sensitive to basic stimulus characteristics (e.g. odor concentration or intensity), whereas LPC amplitude and latency are thought to reflect higher-order evaluative and integrative processing of chemosensory information (Lundström et al., 2006; Pause et al., 1996).

We designed a protocol with four stimulation conditions: a pure trigeminal stimulus (PT; CO2), a pure olfactory stimulus (PO; PEA), an ipsilateral olfactory trigeminal co-stimulation (IOT; CO2 and PEA in the same nostril), and a contralateral olfactory trigeminal co-stimulation (COT; CO2 in one nostril and PEA in the other). In each condition, participants performed a localization task to assess trigeminal sensitivity. It is based on the principle that only trigeminal – but not purely olfactory – stimuli can be spatially discriminated across the two nostrils (Kobal et al., 1989). Behaviorally, we expected (1) PO to be poorly localizable, whereas PT to be reliably localizable; (2) IOT, but no COT, to enhance localization performance relative to PT. In analogy, we expected electrophysiological responses to reflect this pattern: (3) PT to elicit larger and earlier ERP responses than PO; (4) IOT, but not COT, to produce larger amplitudes and shorter latencies than PT; (5) ERP responses to relate to localization performance in trigeminal conditions (PT, IOT, COT), but not in PO.

## Materials and methods

The study was conducted in accordance with the Declaration of Helsinki and approved by the Research Ethics Committee for studies involving humans at the Université du Québec à Trois-Rivières (CER-23-297-07.01). Written informed consent was obtained from all participants prior to participation.

### Participants

A total of 61 participants aged 60 years and older were recruited. Inclusion criteria required (a) unimpaired olfaction, as assessed with the Sniffin’Sticks test battery (Burghart, Wedel, Germany) using age-group adjusted normative values (Oleszkiewicz et al., 2019), (b) unimpaired cognition, as assessed with the Montreal Cognitive Assessment (MoCA; Nasreddine et al., 2005; cut-off ≥ 26), (c) unimpaired episodic memory, as assessed with the Rey Auditory Verbal Learning Test (RAVLT; Lavoie et al., 2018; cut-off Z-score below -1.5 standard deviations for all recall and recognition indices), (d) no clinically relevant depressive symptoms, as assessed by the Geriatric Depression Scale (GDS; Montorio & Izal, 1996; cut-off > 11), as well as (e) no clinically relevant anxiety as assessed by the Geriatric Anxiety Inventory (GAI; Champagne et al., 2018; Pachana et al., 2007; cut-off > 9). This screening battery was administered during a first session.

Based on these criteria, 50 healthy older adults were retained and completed the study (age: 66.7 ± 4.8 years; 34 women). Following EEG pre-processing, participants with fewer than 10 valid trials in at least one of the four conditions were excluded from statistical analyses (Rombaux et al., 2006), resulting in a final sample of 44 participants (age: 66.3 ± 4.6 years; 29 women).

### EEG Recording Set-Up

During a second session, we recorded CSERP using a 32-channel ActiCap linked to an actiCHamp amplifier (BrainVision Products, Montreal, Canada). We positioned the ActiCap according to the international 10-20 system (Klem et al., 1999). We placed reference electrodes on the mastoids and positioned two additional electrodes below the right eye and above the left eye. To optimize electrode–scalp contact, 0.2–0.3 mL of SuperVisc gel (BrainVision Products, Montreal, Canada) was injected into each electrode site using a beveled syringe. Electrode impedances were kept below 10 kΩ throughout the experiment, and data were sampled at 500 Hz.

### Stimulus Delivery and Experimental Conditions

To deliver the chemosensory stimuli, we used a modular olfactometer (OL023; Burghart, Vedel, Germany), which provides a constant airflow of 8 L/min to the participants’ nostrils. The air was humidified to approximately 60% and heated to 36.5°C to prevent irritation (Kobal, 1985). We used two stimuli: (1) carbon dioxide (CO2, intensity: 45%), a pure trigeminal stimulus, and (2) phenyl ethanol (PEA; rose odor, intensity: 40%), a pure olfactory stimulus. Stimuli were delivered under four conditions in a partially randomized order: PT and PO first (order counterbalanced), followed by COT and IOT stimulation (order counterbalanced). Except for the COT condition, when one nostril received a stimulus (CO2 and/or PEA), the other nostril was exposed to odorless air.

Each stimulus was delivered for 200 ms, with an interstimulus interval of 28–30 s to prevent habituation. To maintain vigilance between the stimulation, participants performed a tracking task: using a joystick, they had to keep a circle inside a moving square on a computer screen (Hummel & Kobal, 2001). Rain sound was played through headphones to mask potential auditory cues, e.g., switching clicks of the olfactometer. Additionally, participants were instructed to reduce nasal airflow as much as possible during the experiment.

### Stimulus Localization Task

During the CSERP recording, participants performed a stimulus localization task (Kobal et al., 1989). Specifically, we asked them to indicate which nostril received the stimulus by clicking on a left or right arrow using a hand-held mouse. The task comprised 40 trials, with half of the trials presented to each side in a randomized order. Not all participants completed all trials. To assess participants’ ability to accurately localize stimuli independent of response bias, we calculated the *sensitivity* index d’ (Signal Detection Theory) from the localization performance on trials for which participants provided a response (Hautus et al., 2021).

### EEG Pre-Processing

EEG data were pre-processed using Brainstorm (Tadel et al., 2011), an open-source software package available under the GNU public license (http://neuroimage.usc.edu/brainstorm). As an initial quality control step, power spectrum density (PSD) analyses were computed on all channels of the raw EEG data to identify electrodes exhibiting excessive noise or artifacts. Electrodes consistently contaminated by ocular or frontal artifacts – most notably Fp1 and Fp2 – were excluded from further analyses. A band-pass filter (0.01-30 Hz) was then applied to the raw signals. Independent component analysis (ICA) was subsequently conducted on the filtered data to identify and remove stereotypical artifacts (e.g., eye blinks, heartbeats). Components corresponding to these artifacts were identified by visual inspection of their topographies and time courses and were removed from the data.

### EEG Data Analysis

Electrode clusters and time windows were selected based on recommendations from the literature (Arpaia et al., 2022; Huart et al., 2012; Rombaux et al., 2012). Specifically, the following electrodes were included for analysis: Cz, FC1, FC2, Fz, F3 and F4 for the N1 component, and Cz, CP1, CP2, Pz, P3 and P4 for the LPC component. Using Brainstorm, peak amplitudes and latencies of the N1 (time window: 270-500 ms) and LPC (time window: 460-860 ms) components were extracted at these sites as the minimum (N1) and maximum (LPC) values within the specified time windows. These values were subsequently averaged, yielding one mean amplitude and one mean latency per participant within each condition (PO, PT, COT, IOT) and for each component (N1/LPC).

Outliers were identified and removed prior to the analyses; in total, two condition-level values were excluded (Table 1). Finally, for illustration purposes only, grand-average waveforms were computed at electrode Cz by averaging across participants for each condition, resulting in a single representative waveform per condition (Figure 1).

**Figure 1.**
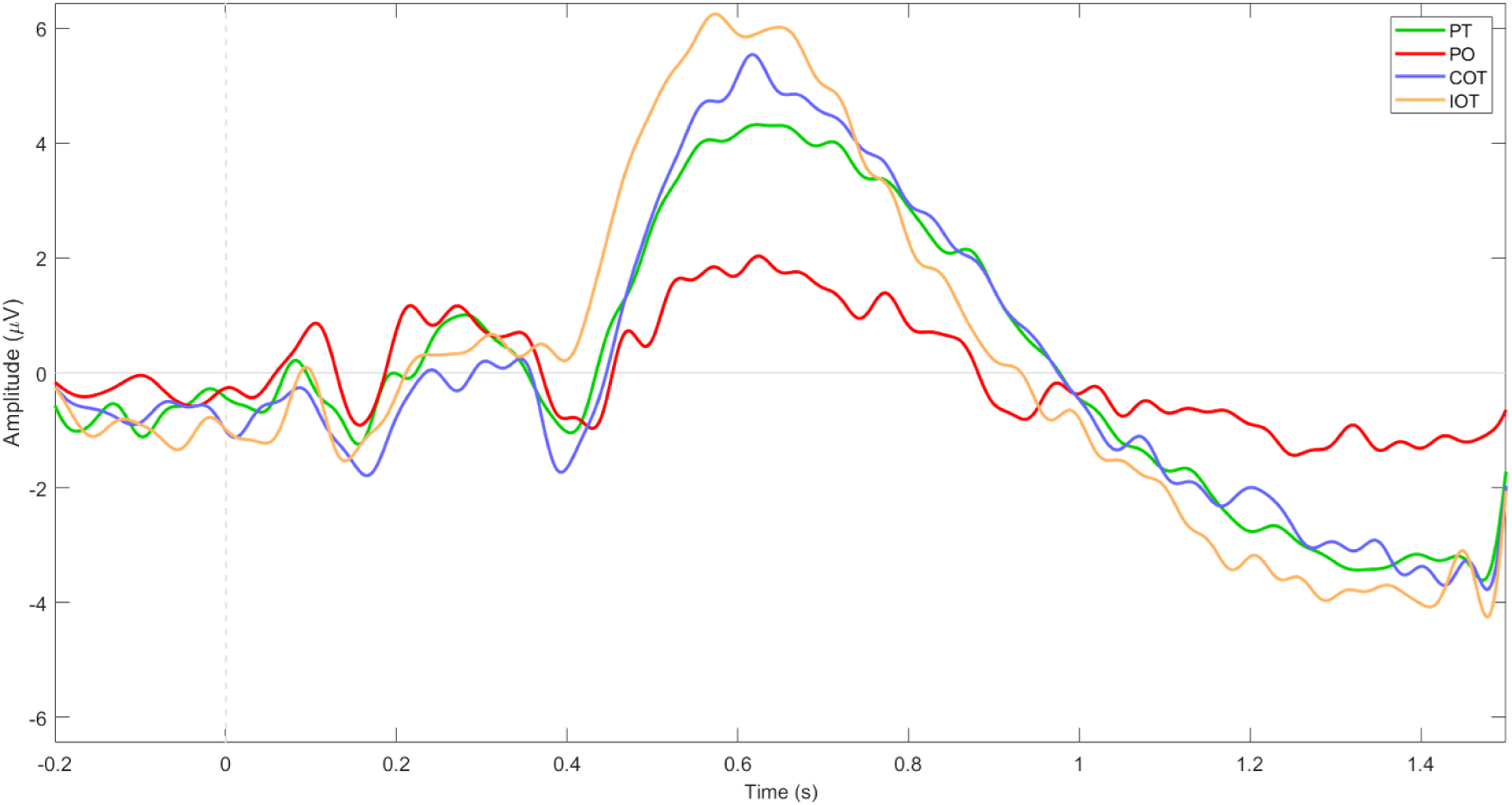
Grand-average waveforms at electrode Cz for each condition. Waveforms represent the average across participants for PT (green), PO (red), COT (blue), and IOT (orange). The dashed vertical line at 0 s indicates stimulus onset (duration: 200 ms). The N1 (270-500 ms) and LPC (460-860 ms) components can be visually identified within their respective time windows. Amplitude is expressed in microvolts (μV) and time in seconds (s). **Alt text**: Line graph of grand-average CSERP waveforms at electrode Cz for PT, PO, COT, and IOT conditions. Trigeminal-containing conditions show larger positive responses than PO, with IOT reaching the highest amplitude around 0.6 s.

**Table 1.**
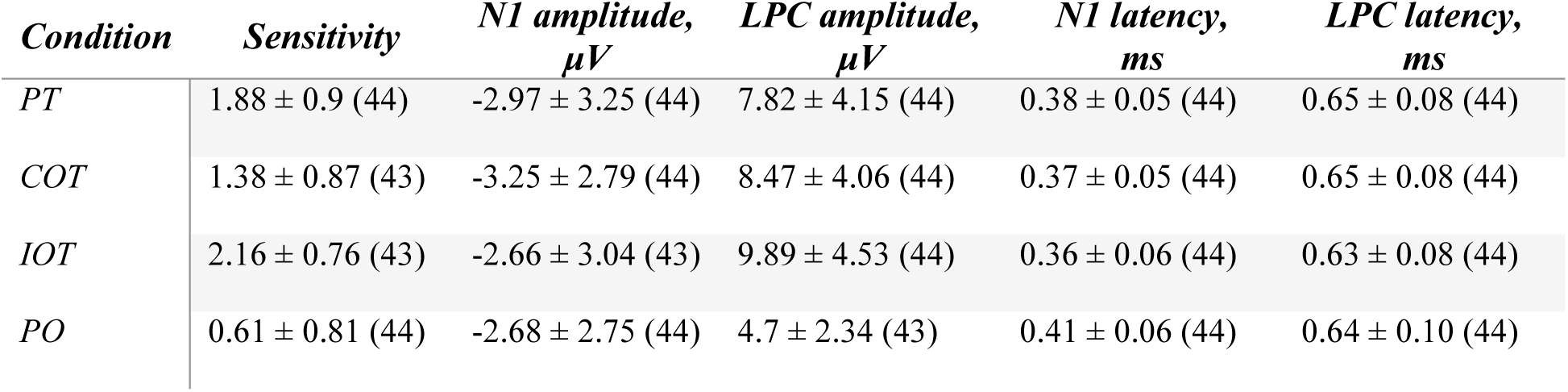
Mean ± SD (n) for sensitivity and CSERP measures by condition.

### Statistical Analysis

All statistical analyses were performed using R Studio (version 2025.09.2). For behavioral data and CSERP components, linear mixed-effects models were applied to examine the effects of stimulus type on each dependent variable. The dependent variables included the *sensitivity* (d’) and the *amplitudes* and *latencies* of the main CSERP components (*N1*, *LPC*). We included *stimulus* (four levels: PO, PT, COT, IOT) as a within-subject factor, and *PO sensitivity* as a covariate to account for potential trigeminal activation of PO. We initially included *age* and *sex* as covariates but then removed them from the final models as they were not significant predictors. When we found significant main effects, we performed Holm-corrected post hoc pairwise comparisons.

We computed partial Spearman correlations separately for each condition to examine associations between *LPC amplitudes* and *sensitivity*, controlling for *age* and *sex*. We corrected the resulting p-values for multiple comparisons across the four conditions using false discovery rate (FDR) correction.

For all analyses, we set statistical significance at p < 0.05.

## Results

### Behavioral results: trigeminal localization test

We observed a significant effect of *stimulus* on the ability to localize stimuli, as expressed by *sensitivity* d’ (F(3, 127.61) = 44.66, *p* < 0.001). Post hoc comparisons showed significant differences between all conditions. Specifically, *sensitivity* was significantly higher in IOT than in PT (*p* < 0.05), COT (*p* < 0.001) and PO (*p <* 0.001), significantly higher in PT than in COT (*p* < 0.01) and PO (*p* < 0.001), and significantly higher in COT than in PO (*p* < 0.001) (Figure 2).

**Figure 2.**
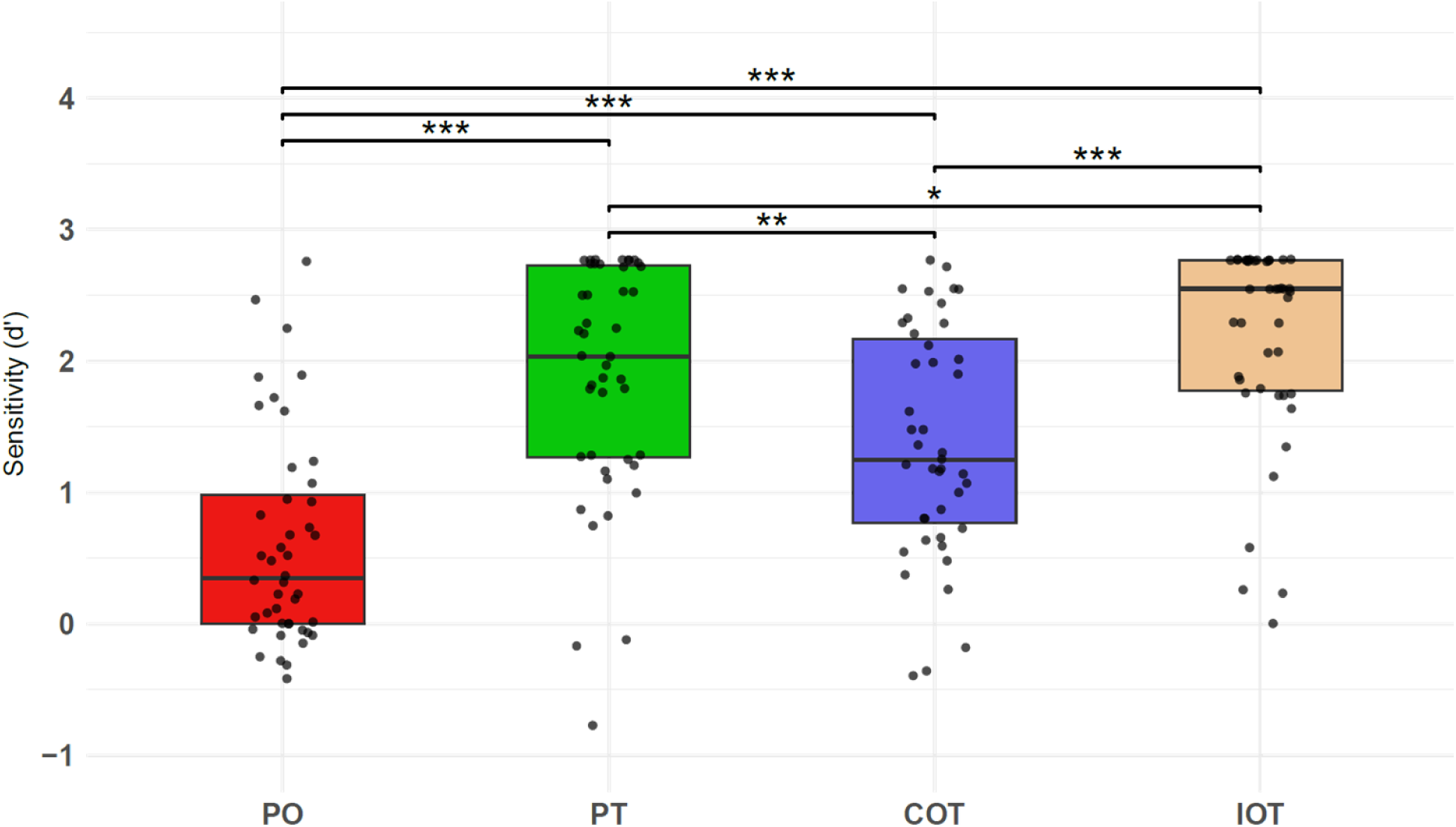
Stimulus localization sensitivity (d’) across conditions. Boxplots represent the distribution of sensitivity scores for each condition: PO (red), PT (green), COT (blue), and IOT (orange). Individual data points are shown as jittered dots. * p<0.05; ** p<0.01; *** p<0.001. **Alt text**: Boxplots show stimulus localization sensitivity scores across PO, PT, COT and IOT conditions. Sensitivity is lowest for PO and highest for IOT, with significant pairwise differences indicated by asterisks above the boxplots. Individual participant data points are shown as jittered dots.

### ERP results: amplitude and latency

We did not observe a significant effect of *stimulus* (F(3, 8.87) = 0.7, p = 0.58), *PO sensitivity* (F(1, 7.32) = 0.1; *p* = 0.76) or interaction between *stimulus*\**PO sensitivity* (F(3, 8.87) = 0.59; *p* = 0.64) on *N1 amplitude*.

In contrast, we observed a significant main effect of *stimulus* on *LPC amplitude* (F(3, 10.52) = 24.33; *p* < 0.001), but no effect of *PO sensitivity* (F(1, 8.43) = 3.12; *p* = 0.11), nor an interaction *stimulus*PO sensitivity* (F(3, 10.52) = 0.75; p = 0.55). Post-hoc comparisons revealed greater amplitudes for all conditions containing a trigeminal stimulus than the PO condition (p < 0.001). Further, IOT elicited significantly larger amplitudes than both PT (*p* < 0.01) and COT (*p* < 0.05), whereas no significant difference was observed between PT and COT (*p* = 0.28) (Figure 3).

**Figure 3.**
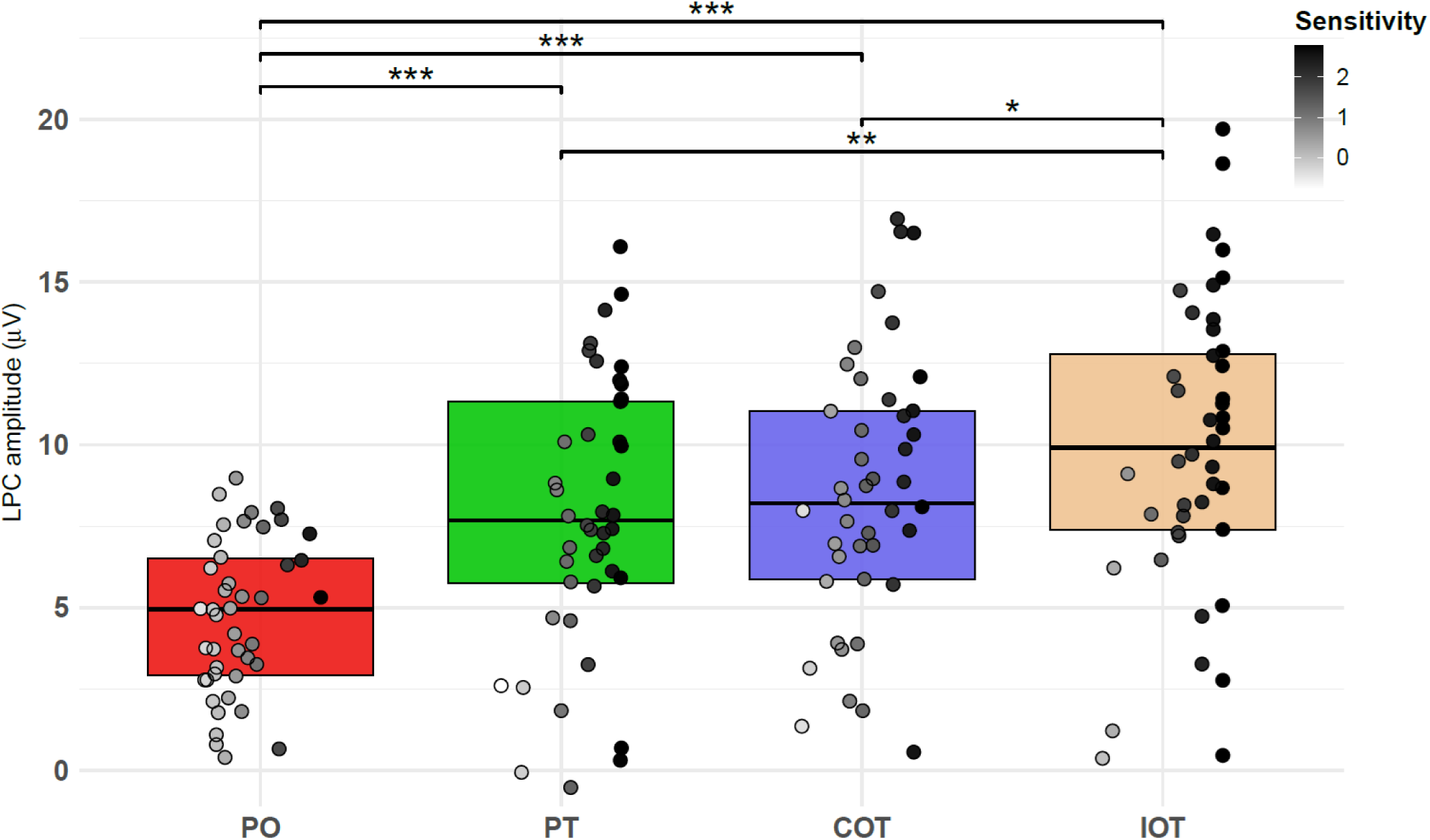
LPC amplitude (μV) across conditions. Boxplots represent LPC amplitudes for each condition: PO (red), PT (green), COT (blue), and IOT (orange). * p<0.05; ** p<0.01; *** p<0.001. **Alt text**: Boxplots show LPC amplitudes across PO, PT, COT, and IOT conditions. LPC amplitude is lowest for PO and highest for IOT, with significant pairwise differences indicated by asterisks above the boxplots. Individual data points are shown as dots, with shading representing stimulus localization sensitivity (darker dots indicating higher sensitivity scores).

We further observed a significant main effect of *stimulus* on *N1 latency* (F(3, 126) = 10.34; *p* < 0.001), while we did not see any effect of *PO sensitivity* (F(1, 42) = 1.8; *p* = 0.19) or an interaction

*stimulus*PO sensitivity* (F(3, 126) = 0.21; *p* = 0.89). Post hoc comparisons showed that *N1 latency* was significantly longer in PO compared to PT (*p* < 0.05), COT (*p* < 0.001), and IOT (*p* < 0.001), but no difference between the three conditions with trigeminal stimulation (Figure 4). In contrast, we did not observe a significant effect of *stimulus* (F(3, 126) = 0.6, p = 0.61), *PO sensitivity* (F(1, 42) = 3.58; *p* = 0.07) or interaction between *stimulus*\**PO sensitivity* (F(3, 126) = 0.78; *p* = 0.5) on *LPC latency*.

**Figure 4.**
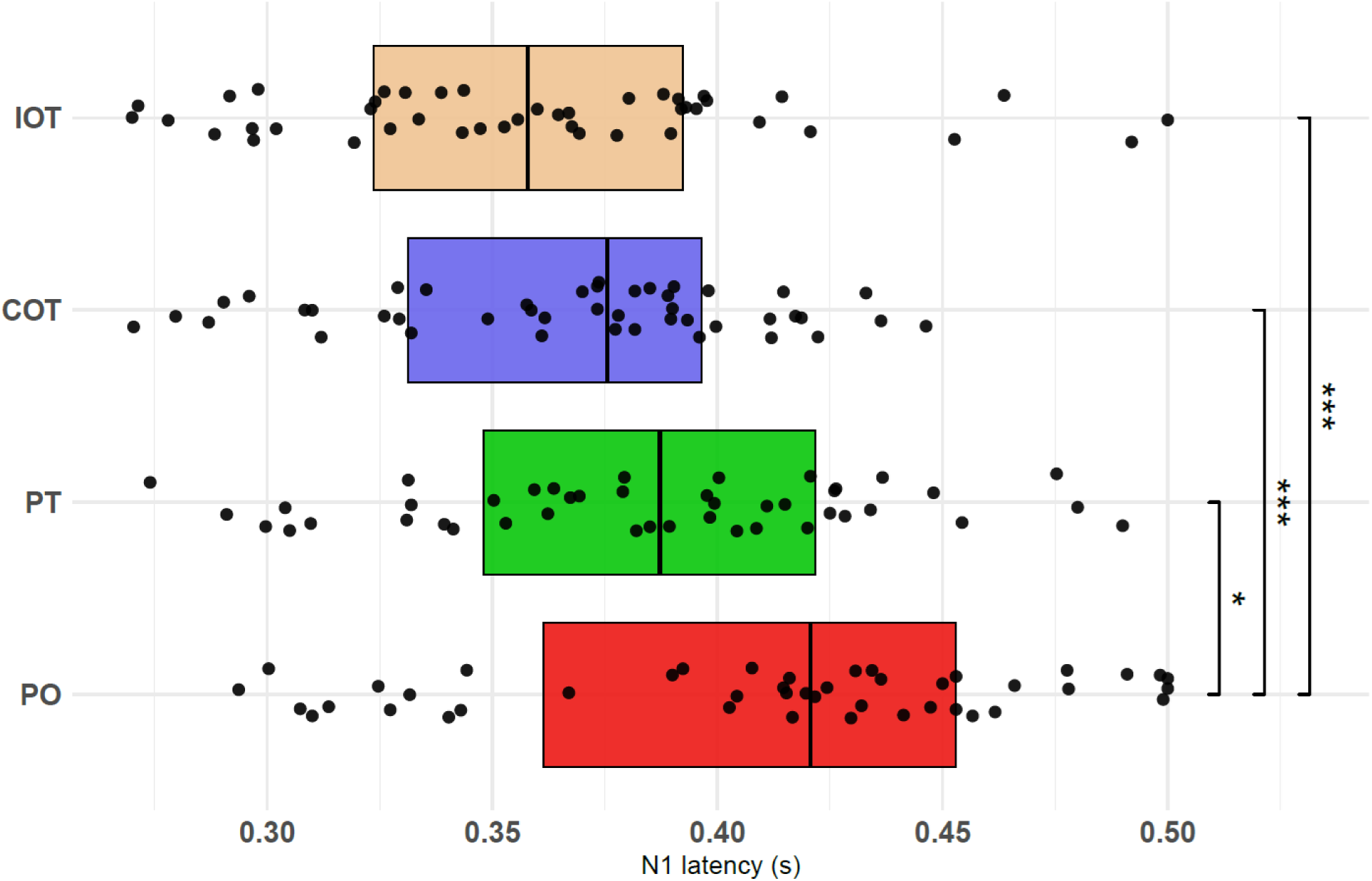
N1 latency (s) across conditions. Boxplots represent N1 latencies for each condition: PO (red), PT (green), COT (blue), and IOT (orange). Individual data points are shown as jittered dots. * p<0.05; ** p<0.01; *** p<0.001. **Alt text**: Horizontal boxplots show N1 latencies across PO, PT, COT, and IOT conditions. PO shows longer N1 latencies than the trigeminal-containing conditions, with significant pairwise differences indicated by asterisks on the right side of the plot. Individual data points are shown as jittered dots.

### Relationship between trigeminal localization sensitivity and LPC amplitude

We observed significant positive correlations between *LPC amplitudes* and *sensitivity* across conditions: PT (ρp = 0.47, q < 0.01, n = 44), COT (ρp = 0.41, q < 0.05, n = 43), IOT (ρp = 0.37, q < 0.05, n = 43), as well as PO (ρp = 0.31, q = 0.0459, n = 43) (Figure 3).

## Discussion

The purpose of this study was to characterize olfactory-trigeminal interactions in healthy older adults by comparing CSERP responses under ipsilateral and contralateral olfactory-trigeminal co-stimulation, to determine the relative contributions of peripheral and central mechanisms. The main finding was that ipsilateral, but not contralateral olfactory-trigeminal co-stimulation, improved trigeminal localization performance and induced larger LPC amplitudes.

In line with our behavioral hypotheses, we found that only ipsilateral olfactory-trigeminal co-stimulation (as in the IOT condition), but not contralateral olfactory-trigeminal co-stimulation (as in the COT condition), enhanced participants’ ability to localize a trigeminal stimulus, indicating that performance depends on the nostril-of-delivery configuration. These findings are consistent with previous studies (Karunanayaka et al., 2020, 2024; Tremblay & Frasnelli, 2018). However, in those studies, trigeminal stimuli were either mixed olfactory-trigeminal compounds (e.g., eucalyptol, mustard oil) or mechanical stimuli (air-puffs), whereas we used CO2, a pure chemosensory trigeminal stimulus, making the interpretation more straightforward. In addition, unlike the earlier reports, we observed significantly stronger responses in PT than COT. One possible explanation is that a subset of participants was able to localize the PO above chance level, suggesting a potential trigeminal contribution. Consequently, in the COT condition, some participants may have perceived trigeminal input from both nostrils simultaneously, potentially creating bilateral ambiguity that may have hindered correct localization of the CO2 stimulus, thereby reducing performance relative to PT.

However, additional analyses indicated that the ability to localize PO did not influence our main findings, as neither PO sensitivity nor its interaction with stimulus reached significance. Nevertheless, these observations highlight the need for caution when using so-called “pure” olfactory stimuli. Although PEA is widely considered a pure olfactory stimulus at low concentrations, it may still induce trigeminal activation in some individuals (Frasnelli et al., 2011). There is currently no clear consensus in the literature regarding the concentration range at which PEA may begin to recruit the intranasal trigeminal system. Moreover, trigeminal activation does not depend solely on concentration, but also on the total amount of stimulus delivered, as both stimulus duration and flow also influence irritancy onset (Cometto-Muñiz & Cain, 1992). This is particularly relevant given that previous CSERP studies in young participants used PEA as a pure olfactory stimulus at concentrations ranging from approximately 10% to over 40% (Bensafi et al., 2007; Flohr et al., 2015; Fortier-Lebel et al., 2023; Sabiniewicz et al., 2024; Welge-Lüssen et al., 2003).

The ERP results indicate that PT was associated with shorter N1 latencies and larger LPC amplitudes than PO. The faster N1 latencies for PT are consistent with evidence that this component is primarily driven by exogenous stimulus properties such as concentration, and trigeminal potency (Nordin et al., 2005; Pause et al., 1996, 1997; Pause & Krauel, 2000). Previous studies, however, have reported mixed findings, with some indicating faster olfactory processing (Geisler & Murphy, 2000), and others showing shorter trigeminal latencies (Hummel & Kobal, 1992; Iannilli et al., 2013). The larger LPC amplitudes observed for PT compared to PO are consistent with previous findings (Flohr et al., 2015).

Regarding CSERP amplitudes, our results suggest that in healthy older adults, olfactory-trigeminal interactions emerge mainly at a later stage of processing, as reflected by enhanced LPC responses rather than by early N1 modulation. While N1 amplitudes did not differ across conditions, IOT elicited significantly higher LPC amplitudes than both PT and COT. Together with the behavioral results, this pattern is consistent with the possibility of a peripheral contribution to olfactory-trigeminal interactions. At the peripheral level, the trigeminal nerve innervates the olfactory mucosa (Brand, 2006), allowing potential interactions between the two systems. Supporting this, a recent study showed that IOT enhances the amplitudes of the Negative Mucosa Potential (NMP), an electrophysiological measure of trigeminal activity in the respiratory mucosa (Kobal, 1985), compared to PT (Mai, Burghardt, et al., 2025). Evidence from rodents further suggests that trigeminal stimulation can trigger the release of neuropeptides such as calcitonin-gene-related peptide (CGRP) and neuromodulators like ATP from trigeminal sensory fibers, which may inhibit excitatory responses in olfactory receptor neurons (Daiber et al., 2013; Genovese et al., 2023). However, an optogenetic study did not find clear evidence for cross-modal interaction between olfactory and trigeminal structures within the nasal cavity (Maurer et al., 2019). Taken together, these results may suggest that IOT strengthens trigeminal input while partly suppressing olfactory input at the peripheral level. However, the existence, extent, and mechanisms of such interactions remain insufficiently understood and insufficiently established in humans (Migneault-Bouchard et al., 2025).

The absence of a significant difference in LPC amplitudes between PT and COT suggests that any COT effect was modest in the present paradigm. This may be partly explained by a violation of the spatial rule of multisensory integration, which holds that integration is enhanced when stimuli originate from the same spatial location (Spence, 2013). In COT, olfactory and trigeminal stimuli are delivered to opposite nostrils, reducing spatial congruency and potentially limiting integration (Karunanayaka et al., 2024). However, this does not imply the absence of central olfactory-trigeminal interactions (Cain & Murphy, 1980). Although the stimuli are anatomically separated at the peripheral level, olfactory information is initially processed ipsilaterally but can rapidly engage bilateral cortical networks via interhemispheric connections (e.g., anterior commissure) (Dalal et al., 2020; Dikeçligil et al., 2023; Lapointe et al., 2025). Consistent with this, single-nostril olfactory stimulation has been shown to evoke bilateral cortical responses (Ekanayake et al., 2024). In contrast, trigeminal inputs reach the cortex through thalamic relays, but are also represented bilaterally, meaning that central convergence of the two systems remains anatomically and functionally plausible even under COT stimulation. The present findings do not provide strong evidence that central interaction mechanisms substantially contributed to LPC responses under the current experimental conditions. One possibility is that COT interactions were not fully captured within the LPC time window examined here. Indeed, olfactory and trigeminal inputs differ in their spatiotemporal cortical processing (Iannilli et al., 2013), and COT processing may involve additional interhemispheric coordination compared to IOT (Dalal et al., 2020). Consequently, COT may follow a slower or less temporally synchronized integration dynamic than IOT, which could account for the absence of robust LPC differences observed in the present study (Huart et al., 2012).

Although how the olfactory and trigeminal systems integrate bilateral sensory inputs remains poorly understood, and the relative contributions of peripheral and central mechanisms remain to be fully clarified, the correlations between LPC amplitudes and behavioral sensitivity suggest that LPC amplitudes reflect functionally meaningful aspects of trigeminal processing. In this context it is worthy to point out that we observed a weak, yet significant, correlation in the PO condition, which may reflect residual trigeminal contributions to PEA perception in some participants, consistent with the above-chance localization performance observed in this condition. Taken together, the correspondence between electrophysiological and behavioral measures aligns with previous work reporting a similar link between CSERP and trigeminal sensitivity (Stuck et al., 2006).

Several limitations should be considered when interpreting the present findings. First, although PEA is typically regarded as a pure olfactory stimulus, some participants were able to localize it above chance. Future studies could address this by using other pure odorants such as vanillin or H2S. Second, although the ipsilateral and contralateral design aimed to dissociate peripheral and central mechanisms, CSERP do not allow for direct assessment of peripheral activation. It would therefore be valuable to combine CSERP with complementary peripheral measures such as NMP and the electro-olfactogram (EOG; Lapid et al., 2009) recorded from the olfactory mucosa. Such multimodal approach would allow a more comprehensive characterization of olfactory-trigeminal interactions and help clarify the respective contributions of peripheral and central mechanisms. Third, the use of predefined ERP time windows may have limited the detection of more distributed or delayed integration processes, particularly for COT stimulation; advanced approaches such as mass univariate analyses with permutation-based correction (Maris & Oostenveld, 2007; Pernet et al., 2015) could provide a more precise characterization of the spatio-temporal dynamics. Beyond these methodological considerations, extending this protocol to clinical populations, such as patients with Parkinson’s disease, would be a promising step toward evaluating the relevance of these markers in pathological contexts involving chemosensory and neurodegenerative dysfunction.

## Authors’ contribution

Legend: C = Conceptualization; M = Methodology; I = Investigation; D = Data curation; A = Analysis; W = Writing – original draft; WE = Writing – review & editing; E = Ethics approval; S = Supervision; FA = Funding acquisition

SB (50 %): C; M; I; D; A; W; WE; E.

OFL (15 %): C; M; I; D; A; WE.

EH (15 %): C; M; I; D; A; WE. KL (5 %): D; A; WE.

JF (15 %): C; M; I; A; WE; E; S; FA.

All authors have read and approved the manuscript.

## Conflicts of interest

The authors declare no conflict of interest.

## Funding

This work was supported by the Fonds de recherche du Québec – Santé [JF: 2024-2025 - CB – 352197], the Fonds de recherche du Québec – Nature et technologies [KL: https://doi.org/10.69777/371202], the Natural Sciences and Engineering Research Council of Canada [JF: 04813-2022; KL: 596163-2024], the Canadian Institutes of Health Research [JF: AFF 173514; OFL: 187458], the Regroupement intersectoriel de recherche en santé de l’Université du Québec [SB], the Quebec Bio-Imaging Network [SB], the Réseau québécois de recherche sur le vieillissement [EH], and the Centre intégré universitaire de santé et de services sociaux du Nord- de-l’Île-de-Montréal [EH].

## Acknowledgements

We thank all the participants for their participation.

## Data Availability

The data underlying this article will be shared in accordance with Ethic Board guidelines on reasonable request to the corresponding author.

## Notes

### Competing Interest Statement

The authors have declared no competing interest.

